# Pharmacokinetic Effects of Combined Exposure to Nicotine and THC via E-Cigarettes in Pregnant Rats

**DOI:** 10.1101/2021.07.07.451537

**Authors:** K. R. Breit, C. G. Rodriguez, S. Hussain, K. Thomas, M. Zeigler, I. Gerasimidis, J. D. Thomas

**Author notes:** **Corresponding author:** Dr. Kristen R. Breit, 125 W Rosedale Ave, Wayne Hall, Room 511, West Chester, PA 19383.

## Abstract

Nicotine and cannabis are two of the most commonly consumed licit and illicit drugs during pregnancy, often consumed together via e-cigarettes. Vaping is assumed to be a safer alternative than traditional routes of consumption, yet the potential consequences of prenatal e-cigarette exposure are largely unknown, particularly when these two drugs are co-consumed. In a novel co-exposure model, pregnant Sprague-Dawley rats received nicotine (36 mg/mL), THC (100 mg/mL), the combination, or the vehicle via e-cigarettes daily from gestational days 5-20, mimicking the first and second human trimesters. Maternal blood samples were collected throughout pregnancy to measure drug and metabolite levels, and core body temperatures before and after exposure were also measured. Pregnant dams exposed to combined nicotine and THC had lower plasma nicotine and cotinine levels than those exposed to nicotine alone; similarly, the combined exposure group also had lower plasma THC and THC metabolite (THC-OH and THC-COOH) levels than those exposed to THC alone. Prenatal nicotine exposure gradually decreased basal core body temperatures each day, with chronic exposure, whereas exposure to THC alone decreased temperatures during the individual sessions. Despite these physiological effects, no changes were observed in food or water intake, weight gain, or basic litter outcomes. These data suggest that combined exposure to nicotine and THC elicits both separate and interactive physiological effects of nicotine and THC on pregnant dams. These data and use of this model can help improve education for pregnant mothers about prenatal e-cigarette use and has important implications for public policy.

**HIGHLIGHTS:**

- Repeated prenatal nicotine exposure via e-cigarettes gradually decreased temperatures
- Prenatal THC exposure via e-cigarettes decreased temperatures during intoxication
- Combined prenatal exposure via e-cigarettes altered plasma drug and metabolite levels
- This co-exposure model elicits separate and interactive effects of nicotine and THC

## 1.1 INTRODUCTION

The potential consequences of nicotine consumption during pregnancy on both the mother and offspring have been extensively studied. Previous research has established that prenatal nicotine exposure can lead to an increased risk of sudden infant death syndrome (SIDS) (Duncan et al., 2009), low birth weight (Ernst et al., 2001; Gatzke-Kopp & Beauchaine, 2007), long-term hypertension, obesity, lung abnormalities, and many other health complications (Bruin et al., 2010). In addition, prenatal nicotine exposure can disrupt development of a number of brain regions and consequent behavioral domains (Jamshed et al., 2020). For example, prenatal nicotine exposure may increase internalizing (Cercone et al., 2015; Cornelius & Day, 2009) and externalizing behavioral problems (Gatzke-Kopp & Beauchaine, 2007; Herrmann et al., 2008; Kahn et al., 2003), hyperactivity (Ernst et al., 2001; Kahn et al., 2003) and ADHD (Banerjee et al., 2007; Biederman, 2005; Herrmann et al., 2008; Linnet et al., 2003; Mick et al., 2002), and can impair learning and memory (Cornelius et al., 2001; Longo et al., 2014), language (Fried & Watkinson, 1990; Fried et al., 1997), and cognitive performance (Derauf et al., 2009; Ernst et al., 2001; Fried & Watkinson, 1988; Fried et al., 1992; Fried & Watkinson, 1990; Herrmann et al., 2008).

Importantly, though, the routes of administration for nicotine consumption have drastically changed. The use of electronic cigarettes (e-cigarettes) has gained extensive popularity over the past several years, even among pregnant women. Recent reports indicate that 5-14% of pregnant women report using e-cigarettes during pregnancy (Cardenas et al., 2019). This is largely in part due to the widespread assumption that e-cigarette use is safer than traditional smoking routes. Among pregnant women with equivalent knowledge about the dangers of traditional smoking during pregnancy, 43% report believing that vaping is a safer alternative (Mark et al., 2015).

At this time, it is unclear how consumption via vaping could affect fetal development, especially since the e-cigarette vehicle (propylene glycol and other chemicals) contains constituents that may, by themselves, exert damaging effects (Strongin, 2019). Unfortunately, research in this area is severely limited, but the research that does exist illustrates that prenatal e-cigarette use induces fetal changes in DNA methylation, decreases birth weight, and increases birth defects (Cardenas et al., 2019). However, no research has examined the long-term effects of prenatal e-cigarette exposure, despite requests from medical professionals (Suter et al., 2015). Given that research has shown that drug consumption via e-cigarettes leads to higher drug and metabolite levels in the blood, including maternal and fetal blood (Young-Wolff et al., 2020), it is possible that drug consumption via e-cigarettes could lead to more severe consequences than previously demonstrated with traditional cigarette use.

Moreover, the design of e-cigarettes allows for multiple substances to be consumed simultaneously, which could be more detrimental than exposure to any drug alone. For example, nicotine and cannabis are more commonly consumed together than cannabis alone among pregnant women (Coleman-Cowger et al., 2017; Gray et al., 2010). On its own, the effects of prenatal cannabis exposure are still not well understood, despite an increase in cannabis use among women of child-bearing age in the U.S. (Brown et al., 2017), including pregnant women (Agrawal et al., 2019), particularly during the first trimester (Volkow et al., 2019). Reports from 2014-2017 estimate that 4.5% of pregnant women had consumed cannabis in the past 30 days, with over 8% of pregnant women in the first trimester consuming cannabis in the past 30 days (Young-Wolff et al., 2020). These numbers have likely continued to rise, as pregnant women may purposefully consume cannabis products to combat nausea and other pregnancy symptoms, believing that gestational cannabis use is safe (Brown et al., 2017). This assumption is indeed unfounded, as more recent research has shown that cannabis use during pregnancy can actually provoke recurrent nausea and vomiting rather than combat it (Kim et al., 2018) and can interact negatively with other required medications (Brown & Winterstein, 2019; Ujváry & Hanuš, 2016).

To date, both clinical and preclinical studies examining cannabis exposure during pregnancy have yielded inconsistent results. Results are largely dependent upon the cannabinoids consumed, timing of exposure, doses, routes of administration, and a host of other factors (Huizink, 2014; Schneider, 2009). Although outcomes vary, some recorded alterations include reduced birth weight (Day et al., 1991; Day & Richardson, 1991; Fergusson et al., 2002; Fried & O’connell, 1987; Huizink, 2014; Hurd et al., 2005) and altered emotional, behavioral, and cognitive development (Huizink, 2014). Importantly, the potency of the primary psychoactive component of cannabis, THC, has risen drastically in the past few decades from 3.4% in 1993 (Chandra et al., 2019; Mehmedic et al., 2010) to the average potency of 17.1% (Chandra et al., 2019). Levels are even higher among synthetic cannabinoid preparations (e.g. Spice) compared to plant cannabis (Botticelli, 2016; Subbanna et al., 2013).

Moreover, similar to nicotine consumption trends, the use of e-cigarettes to consume cannabis constituents has also gained popularity. Still, we know very little about the potential effects of vaporized cannabis consumption. Thus, the results from research from previous years may not be applicable to the levels and routes of cannabis consumption that are currently used today. Several ongoing clinical prospective studies are addressing these questions to better understand the potential consequences of prenatal cannabis consumption in current times, including the ABCD study (Paul et al., 2019); however, the results of these studies will not be known for years to come.

Given that nicotine and cannabis can be easily consumed simultaneously using e-cigarette tanks, it is important to examine the possible interactive effects of co-consumption of these drugs. This is especially critical given that 40% of pregnancies in the U.S. are unplanned (Sedgh et al., 2014), and co-consumption is highest among child-bearing age groups. For example, one study of undergraduate midwestern college students found that 77.9% of those who used e-cigarettes to consume nicotine also used e-cigarettes to consume cannabis products (Kenne et al., 2017). Importantly, when these two drugs are co-consumed via e-cigarettes, it may lead to additional toxicant exposure compared to use of either drug alone (Smith et al., 2019). Thus, it is vital to examine the possible interactions of nicotine and cannabis constituents on maternal physiology, so that we can explore whether combined use could be more damaging to a developing fetus.

The current study sought to develop a model of combined prenatal exposure to nicotine and THC via e-cigarettes using a rodent model. Studies using rodent models can help provide quick, accessible information regarding the potential consequences of prenatal nicotine and THC exposure at the levels and routes currently being consumed. Pregnant Sprague-Dawley rats were exposed to nicotine, THC, the combination, or the vehicle during a period equivalent to the human first and second trimesters. Throughout the e-cigarette exposure period, maternal body weights, food intake, and water intake were recorded. In addition, core body temperatures and blood drug and metabolite levels were collected to determine potential pharmacokinetic effects in pregnant subjects. Following birth, the length of gestation, litter characteristics, and early developmental milestones were also recorded to monitor basic litter outcomes following prenatal e-cigarette exposure.

## 2.1 MATERIALS AND METHODS

### 2.1.1 Subjects

This study developed a prenatal co-exposure model of nicotine and THC via e-cigarettes in pregnant rats (Figure 1). All procedures included in this study were approved by the San Diego State University (SDSU) Institutional Animal Care and Use Committee (IACUC) and are in accordance with the National Institute of Health’s *Guide for the Care and Use of Laboratory Animals*. Naïve female Sprague-Dawley rats were obtained from Charles River Laboratories (Hollister, CA) on postnatal day (PD) 60 and allowed to acclimate for at least two weeks prior to any handling or procedures in the animal care facilities at the Center for Behavioral Teratology (CBT) at SDSU.

**Figure 1.**
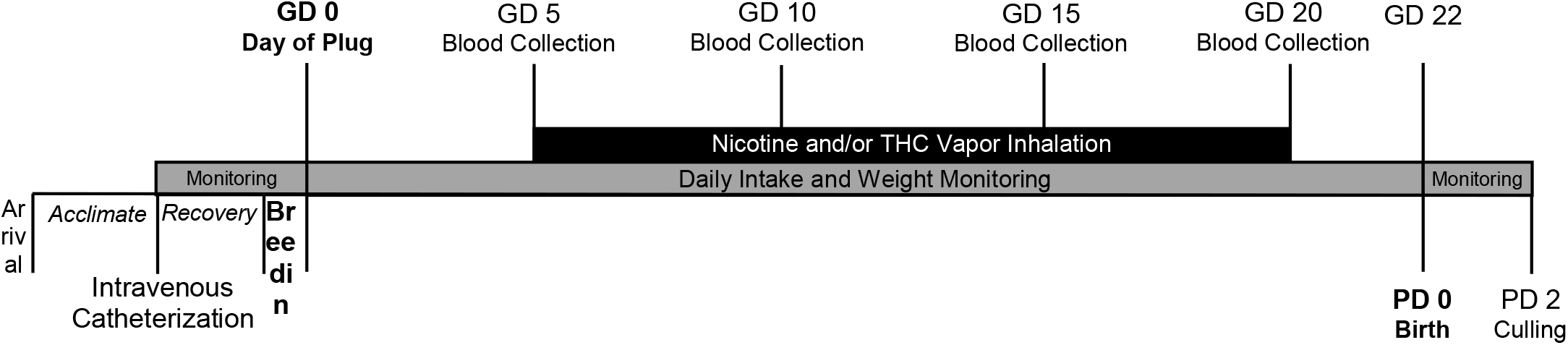
Timeline of study events.

#### 2.1.1.1 Intravenous Catheterization

In order to measure maternal blood levels of each drug and their metabolites, intravenous catheters were surgically implanted prior to pregnancy following a two-week acclimation period, as previously published (Breit et al., 2020). Dams were anesthetized (4% isoflurane) and the catheter was secured under the skin on the back, with the tubing thread subdermally over the right shoulder and implanted into the right jugular vein via small incisions. Tubing was flushed (heparinized bacteriostatic saline) and cuts were sealed (VetBond; 3M). Catheters were then covered with a plastic hood and a metal screw cap to prevent chewing. Dams were administered an antibiotic (Cefazolin, 100 mg/mL; Victor Medical) and a painkiller (Flunixin, 2.5 mg/mL; Bimeda) following surgery for 2 consecutive days post-surgery (subcutaneous injection, 0.001 mL/g).

#### 2.1.1.2 Breeding

Following surgical recovery (~1-2 weeks), dams were paired with a male stud to breed. Breeding pairs were housed in standard Allentown rat cages with raised grid wire floors; a filter paper was placed under the wire floor to catch any seminal plugs. Throughout breeding, pairs had *ad libitum* access to food and water. Each morning, pairs were checked for the presence of a seminal plug, which was designated as gestational day (GD) 0.

### 2.1.2 Prenatal Vapor Inhalation Exposure

On GD 0, pregnant dams were assigned to 1 of 4 prenatal exposure groups: nicotine (36 mg/mL), THC (100 mg/mL), the combination, or the vehicle (propylene glycol). Beginning on GD 0, pregnant dams were monitored daily for gestational weight gain, as well as food and water intake. Dams were weighed prior to any procedures each morning. In addition, food and water intake from the previous day was measured and replenished each morning. Dams were given *ad libitum* access to 200 g of standard pellet lab chow (LabDiet 5001) and 400 mL of water in a graduated water bottle each day.

#### 2.1.2.1 Drugs

THC for e-cigarette vapor inhalation was obtained through the National Institutes of Drug Abuse Drug Supply Program and was prepared as described in (Breit et al., 2020). Powdered nicotine (nicotine hydrogen tartrate salt; Sigma-Aldrich) was added to propylene glycol (Sigma-Aldrich) and vortexed to obtain the required concentration (36 mg/mL).

#### 2.1.2.2 Vapor Inhalation Paradigm

Vaporized drug was administered via commercially available e-cigarette tanks (SMOK V8 X-Baby Q2) in two 4-chamber vapor inhalation apparatuses designed by La Jolla Alcohol Research Inc. (La Jolla, CA, USA). The vapor inhalation system used a sealed cage identical in structure and dimensions as standard rat Allentown cages (Allentown, PA), a computer-controllable adapter to trigger e-cigarette tanks (SMOK V8 X-Baby Q2), and vacuum regulation of the air/vapor flow.

From GD 5-20, pregnant dams were placed in the vapor inhalation chambers for 40 minutes each day; drugs were administered through individual 6-sec puffs, with puffs occurring once every 5 min through steady airflow (2 L/min) for 30 min (7 puffs total). Dams remained in the chambers for an additional 10 min to clear out any residual vapor before removal.

#### 2.1.2.3 Core Body Temperatures

As demonstrated in our lab (Breit et al., 2020) and others (Javadi-Paydar et al., 2018; Nguyen et al., 2016), THC via e-cigarettes lowers core body temperatures in female rats. Maternal body temperatures were measured before and after each vapor exposure session via a rectal thermometer, to verify this expected physiological effect and to examine effects of nicotine exposure on body temperature.

#### 2.1.2.4 Plasma Drug Levels

Blood samples were taken via the intravenous catheters throughout pregnancy to determine drug levels, establish a time curve for each drug, and examine possible pharmacokinetic interactions. On GD 5, 10, 15, and 20, 700 μL of blood were taken at 15-, 30-, 60-, 90-, and 180-min post-vapor exposure. Prior to and after each collection, the dam’s catheter was flushed with heparinized saline (0.2 mL). In the rare case that blood could not be successfully drawn via the catheter, blood was instead taken via tail vein cut. Blood samples were immediately placed in the centrifuge to separate plasma and stored at −80°C until analyses were conducted.

THC, nicotine, and all metabolite levels were analyzed by MZ Biolabs (Tucson, AZ). Methodology for THC and metabolite (THC-OH, THC-COOH) analyses are described in full in (Breit et al., 2020). For nicotine and metabolite (cotinine) analyses, 50 μL of plasma was precipitated using 200 μl HPLC grade acetonitrile containing 10 ng/ml nicotine-D_4_ (Cerilliant N-048-1ML) and cotinine-D_3_ (Cerilliant C-017-1ML) as internal standards. Samples were vortexed vigorously and incubated at 4°C for 30 minutes to precipitate proteins. After a 10-min centrifugation at 4°C, supernatant was transferred to a 96-well plate for analysis using LC-MS2. An 8-point standard curve containing nicotine (Cerilliant N-008-1ML), and cotinine (Cerilliant C-016-1ML) was prepared with concentrations of both analytes ranging from 781 pg/mL to 100 ng/mL. A Thermo Scientific Surveyor HPLC connected to a Thermo Scientific LTQ Velos Pro mass spectrometer was used for separation and quantification of nicotine and cotinine. The Surveyor sample compartment was kept at 6°C and column temperature maintained at 25°C. Analytes were separated using a 4.6 mm × 150 mm HILIC column (Phenomenex 00F-4449-E0) with isocratic flow of 500 μL/min. Isocratic conditions were 5% A, 95% B for 7 min, where A is water containing 25 mM formic acid, 1 mM trifluoroacetic acid and B is acetonitrile. Eluate was analyzed by the LTQ Velos Pro using positive ions defined in the following table. Quantitation was performed using the Quan Browser software from Thermo Scientific. An in-house QC was prepared with 10 ng/mL each of nicotine and cotinine prepared in drug free canine plasma. The linear quantitative range of the assay was from 781 pg/mL to 100 ng/mL.

#### 2.1.2.5 Birth and Litter Monitoring

Gestational lengths were recorded(usually GD 22). On postnatal day (PD) 2, the number of pups born was counted, pups’ sexes were recorded, and each pup was weighed to examine litter size, litter sex ratio, and average pup weight. Litters were pseudorandomly culled to 8 pups (4 females and 4 males, whenever possible) for future behavioral examination. Offspring were monitored daily for eye opening (both eyes fully open), a physical developmental milestone.

### 2.1.3 Statistical Analyses

All data were analyzed using the Statistical Packages for Social Sciences (SPSS, version 26; IBM). Data were analyzed using 2 (Nicotine) x 2 (THC) Analyses of Variance (ANOVAs) with significance levels set at *p* < 0.05. Data analyzed over multiple Days and/or Time points used these within-subject variables as Repeated Measures. Student Newman Keuls post hoc tests (*p* < 0.05) were used when needed. Offspring data additionally used Sex as a between-subject variable. For eye opening, nonparametric analyses were used.

## 3.1 RESULTS

A total of 48 female Sprague-Dawley rats completed the vapor inhalation exposure procedure and gave birth (Nicotine+THC: 12, Nicotine: 12, THC: 11, Vehicle: 13). No complications were observed among pregnant dams; 5 dams were dropped from the study as they were not pregnant (Nicotine+THC: 0, Nicotine: 2, THC: 1, Vehicle: 2).

### 3.1.1 Body Weight Gain

Body weight gain among pregnant dams was analyzed using a 2 (Nicotine) × 2 (THC) ANOVA with Day as a repeated measure. Although all subjects gained weight during gestation (Day: F[21,924] = 1077.00, *p* < 0.001), a 2-way Day*Nicotine interaction was observed (F[21,924] = 1.70, *p* < 0.05). However, there were no main or interactive effects of Nicotine or THC on any individual Day (Figure 2A). Similarly, the percentage of weight gain among dams (GD 0-21) was not significantly altered by prenatal nicotine, THC, or combined exposure during gestation (Figure 2B).

**Figure 2.**
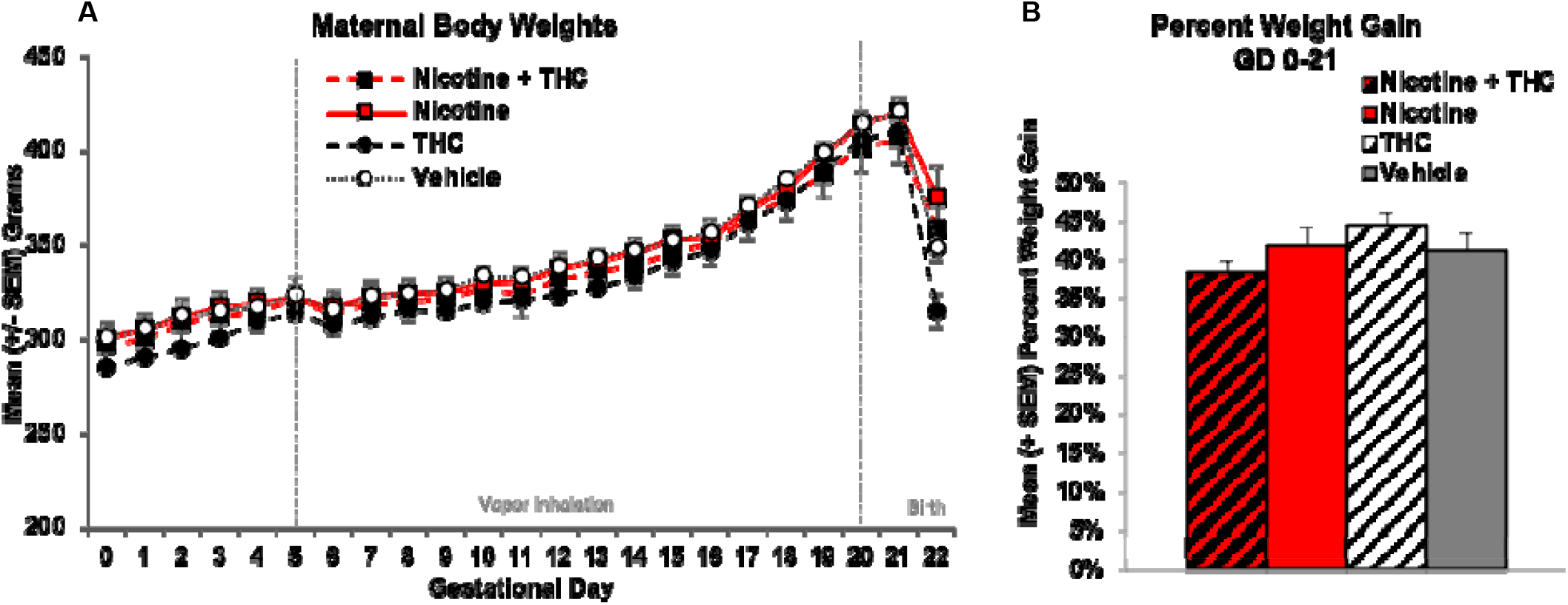
Neither maternal body weights (A) nor pregnancy weight gain (B) during gestation were significantly altered by prenatal exposure to nicotine, THC, or the combination via e-cigarettes.

### 3.1.2 Food and Water Intake

To examine whether prenatal nicotine or THC exposure altered nutritional factors, food and water intake were recorded each morning (for the previous day) prior to vapor exposure. Food and water intake were analyzed separately (1) prior to the vapor inhalation exposure onset (GD 0-4) and (2) during the vapor inhalation period (GD 5-20). All intake data were collapsed over 4-day epochs for simplicity of presentation (GD 0-4, GD 5-8, GD 9-12, GD 13-16, GD 17-20).

Prior to vapor inhalation, dams assigned to receive THC exposure (alone or in combination with nicotine) ate more food at baseline (F[1,44] = 4.11, *p* < 0.05), an artifact of random assignment since they had not yet received any drug exposure. Pregnant dams increased food intake throughout the vapor inhalation period (Day: F[3,132] = 50.45, *p* < 0.001), but neither prenatal nicotine nor THC exposure significantly altered food intake overall, across Days, or on individual Days (Figure 3A). Thus, prenatal exposures did not affect food intake.

**Figure 3.**
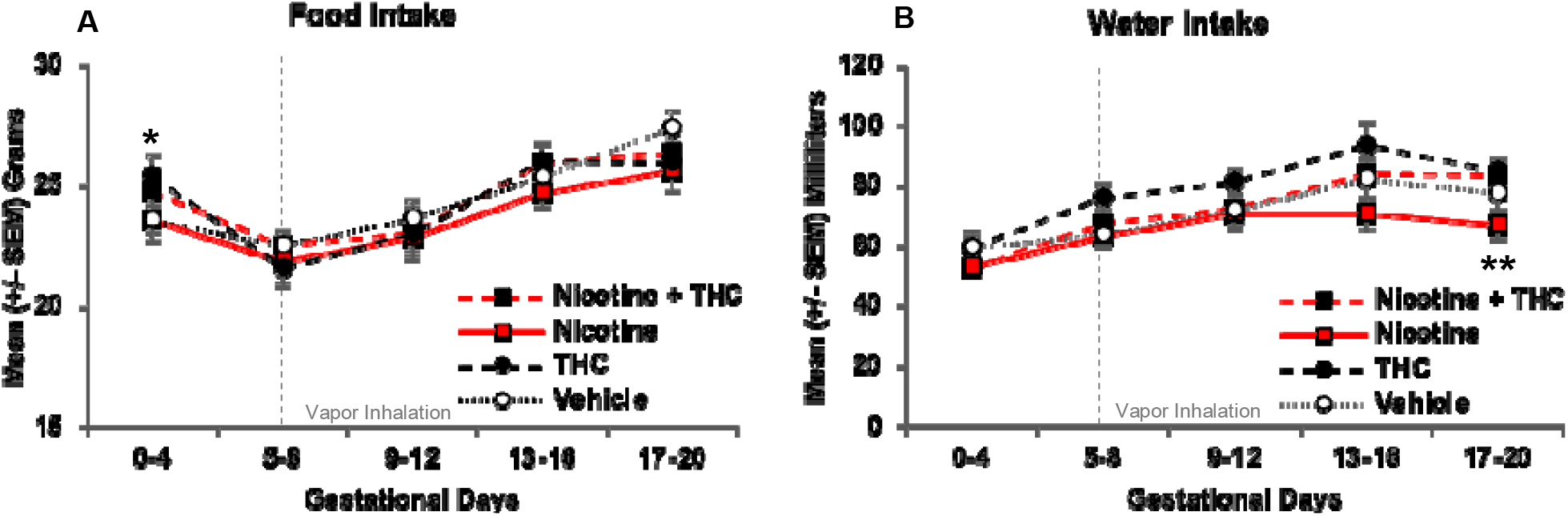
At baseline, pregnant dams assigned to receive THC exposure had higher food intakes, however, no differences among groups were observed throughout the vapor inhalation period (A). During the last days of vapor inhalation, dams exposed to nicotine alone during pregnancy drank less water than dams exposed to THC, although no groups differed significantly from controls (B). * = THC > no THC, *p* < 0.05. ** = Nicotine < Nicotine+THC and THC, *p*’s < 0.05.

At baseline (GD 0-4), no significant differences were seen among groups for water intake. Dams increased water intake throughout the vapor inhalation period (Day: F[3,132] = 23.17, *p* < 0.001); a 3-way interaction of Day*Nicotine*THC approached significance (F[3,132] = 2.38, *p* = 0.07) and there was a main effect of THC (F[1,44] = 4.63, *p* < 0.05). There were no effects of prenatal drug exposure from GD 5-16; however, by GD 17-20, pregnant dams exposed to Nicotine alone drank less water than those exposed to THC or the combination of Nicotine+THC (F[3,44] = 3.19, *p* < 0.05; SNK *p*’s < 0.05). However, water intake did not differ between any prenatal drug-exposed group and Vehicle controls (Figure 3B).

### 3.1.3 Maternal Body Temperature

Maternal core body temperatures were taken before and after vapor exposure each day. Temperature data were collapsed every 4 Days for presentation simplicity (GD 5-8, GD 9-12, GD 13-16, GD 17-20), and analyses used 2 (Nicotine) x 2 (THC) ANOVAs with Day as a repeated measure.

Overall, gestational nicotine exposure via e-cigarettes decreased baseline core body temperatures (F[1,44] = 29.06, *p* < 0.001; Figure 4A). However, core temperatures changed across Days and varied by prenatal exposure group, producing interactions of Days*Nicotine*THC (F[3,132] = 5.10, *p* < 0.01), Days*Nicotine (F[3,132] = 4.59, *p* < 0.01), and Days*THC (F[3,132] = 8.34, *p* < 0.001). On GD 5, prior to any vapor exposure, there were no significant differences in basal body temperatures. During the first half of treatment (GD 6-12), dams exposed to THC alone exhibited higher basal temperatures than all other groups (GD 5-8: F[3,44] = 5.33, *p* < 0.01, SNK *p*’s < 0.05; GD 9-12: F[3,44] = 12.57, *p* < 0.001, SNK *p*’s < 0.05). However, body temperatures of pregnant dams exposed to any of the drugs (combined Nicotine+THC: F[1,11] = 62.35, *p* < 0.001; Nicotine alone: F[1,11] = 37.36, *p* < 0.001; THC alone: F[1,10] = 66.39, *p* < 0.001) gradually decreased over the course of treatment, whereas maternal temperatures of Vehicle controls did not change throughout treatment. By the second half of the exposure period (GD 13-20), dams exposed to Nicotine via e-cigarette, alone or in combination with THC, exhibited significantly lower basal temperatures than those exposed to THC only or the Vehicle (GD 13-16: F[1,44] = 16.62, *p* < 0.001; GD 17-20: F[1,44] = 31.60, *p* < 0.001; Figure 4B).

**Figure 4.**
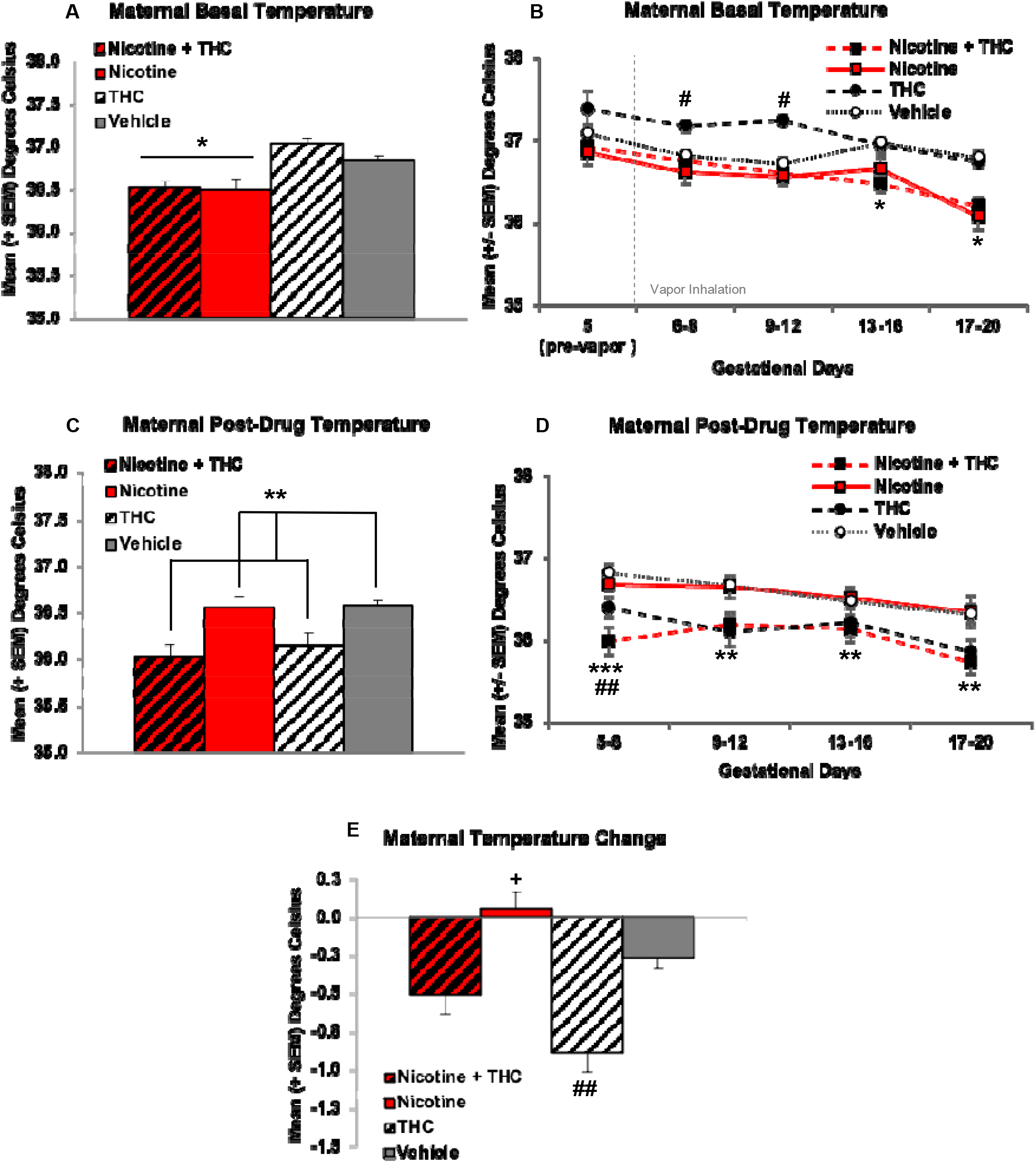
Pregnant dams exposed to nicotine via e-cigarettes had lower basal temperatures across the vapor inhalation exposure period (A). At the beginning of drug exposure, dams exposed to THC had significantly higher basal temperatures, which declined to control levels by mid-pregnancy. Dams exposed to any nicotine maintained lower basal temperatures throughout the latter half of pregnancy (B). In contrast, pregnant dams exposed to THC via e-cigarettes had lower temperatures following vapor inhalation (C). Early in exposure, this effect was driven by the combined exposure group, but a main effect of THC remained throughout the rest of the exposure period (D). When compared against basal body temperatures, pregnant dams exposed to THC had a greater temperature change compared to all other groups (E). * = Nicotine different than no Nicotine, *p*’s < 0.05. # = THC only > all other groups, *p* < 0.01. **= THC < no THC, *p*’s < 0.05. *** = Nicotine+THC < all other groups, *p*’s < 0.001. + = Nicotine only > Vehicle, *p* < 0.05. ## = THC only < Vehicle, *p* < 0.05.

Following intoxication, immediately after drug exposure, dams exposed to THC had lower body temperatures, as expected (F[1,44] = 17.19, *p* < 0.001; Figure 4C). In addition, a Day*Nicotine interaction was also observed (F[3,132] = 2.71, *p* < 0.05). During GD 5-8, dams exposed to combined Nicotine+THC had lower post-drug exposure temperatures than all other groups, whereas those exposed to THC alone had lower body temperatures compared to Vehicle controls, and those exposed to Nicotine alone did not differ from any groups (F3,44] = 9.36, *p* < 0.001; SNK *p*’s < 0.05). However, throughout the rest of the exposure period (GD 9-20), a main effect of THC remained, where dams exposed to any THC had lower ending temperatures than those not exposed to THC (GD 9-12: F[1,44] = 14.22, *p* < 0.001; GD 13-16: F[1,44] = 5.17, *p* < 0.05; GD 17-20: F[1,44[ = 13.10, *p* < 0.01; Figure 4D).

Because of the nicotine-related reductions in baseline body temperature, when examining the temperature change (post-inhalation – pre-inhalation body temperature), dams exposed to THC alone had significantly greater temperature changes and dams exposed to Nicotine alone had significantly smaller temperature changes compared to the Vehicle controls. Dams exposed to combined Nicotine+THC had an intermediate effect, not differing significantly from controls (F[3,44] = 12.83, *p* < 0.001, SNK *p*’s < 0.05; Figure 4E).

### 3.1.4 Blood Level Analyses

All plasma drug and drug metabolite levels were analyzed using 2 (Nicotine) x 2 (THC) ANOVAs with Day and Time as repeated measures. Samples from 3 dams in the Nicotine only group were not analyzed due to technical difficulties. Samples from the Vehicle-exposed subjects were collected, but not analyzed.

#### 3.1.4.1 Nicotine and Metabolite Levels

Dams exposed to the combination of Nicotine+THC via e-cigarettes had lower plasma nicotine levels than those exposed to Nicotine alone (F[1,19] = 37.38, *p* < 0.001; Figure 5). Group differences were most robust close to peak intoxication periods, producing an interaction of Time*THC (F[4,76] = 6.55, *p* < 0.01). Although the Day*Time*THC interaction did not reach statistical significance, plasma nicotine levels of dams exposed to combined Nicotine+THC, but not nicotine alone, gradually increased throughout pregnancy (F[3,33] = 4.99, *p* < 0.01). In fact, by the last day of vapor inhalation (GD 20), nicotine levels of dams exposed to combined Nicotine+THC were more consistent with those exposed to Nicotine alone, differing significantly only at 15 and 90 min (15-min: F[1,19] = 7.45, *p* < 0.05; 90-min: F[1,19] = 4.66, *p* < 0.05; Figure 5D).

**Figure 5.**
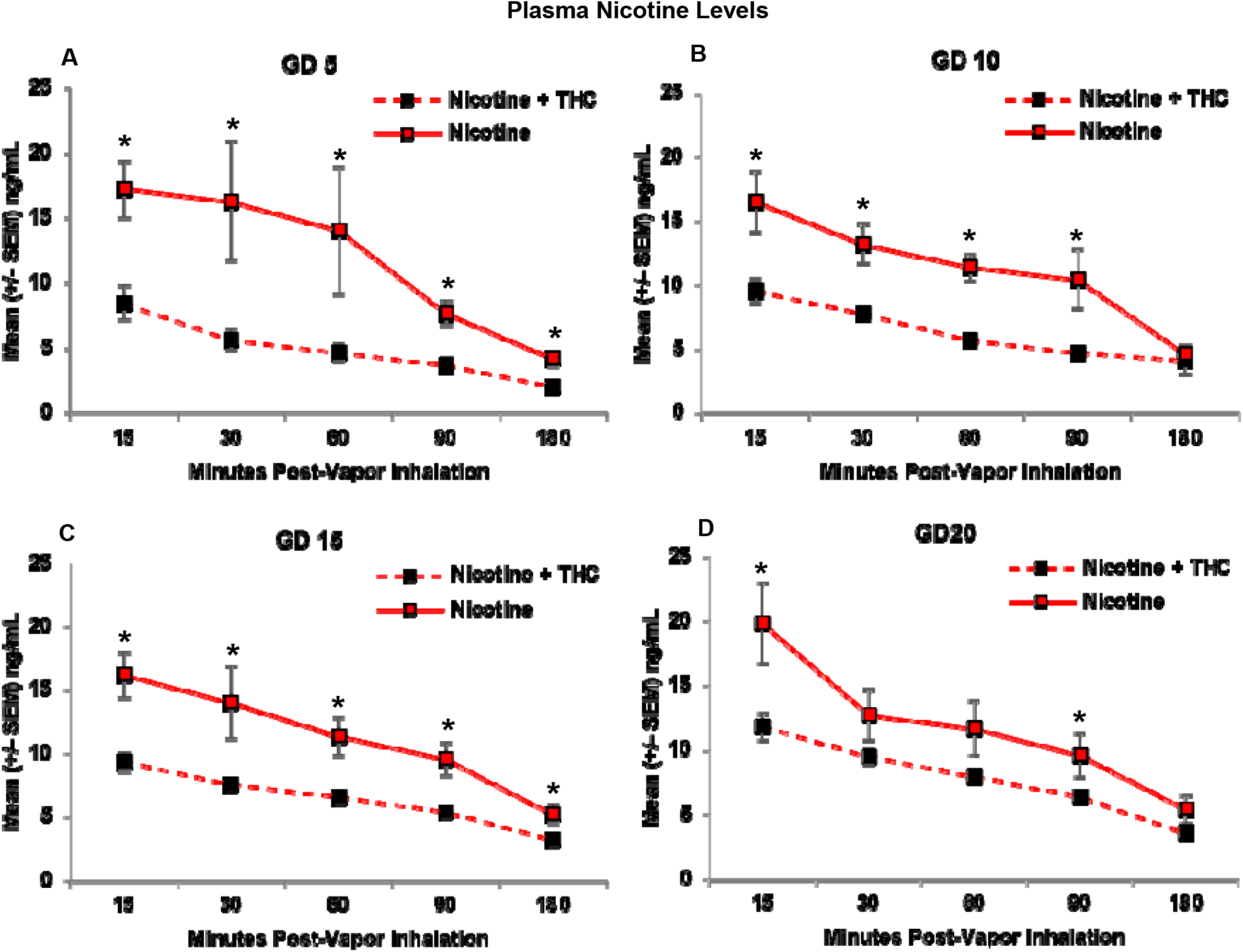
The addition of THC significantly reduced plasma nicotine levels. This effect was seen consistently on GD 5 (A), 10 (B), and 15 (C). By GD 20, this effect was still present, but not as pronounced, as nicotine levels in the combined exposure group gradually increased throughout the vapor inhalation exposure period (D). * = Nicotine+THC < Nicotine, *p* < 0.05.

Consistent with nicotine levels, plasma cotinine levels were also higher in pregnant dams exposed to combined Nicotine+THC compared to those exposed to Nicotine alone (F[1,19] = 40.20, *p* < 0.001), an effect that became less pronounced over days, producing a 3-way interaction of Day*Time*THC (F[12,228] = 3.53, *p* < 0.001). Although both exposure groups showed an expected rise in cotinine levels over Time (*p*’s < 0.001), only dams exposed to combined Nicotine+THC showed an increase in average levels across Days (F[3,33] = 4.99, *p* < 0.01). Importantly, the Nicotine+THC-exposed dams exhibited lower cotinine levels on each Day and Time point (*p*’s < 0.05; Figure 6).

**Figure 6.**
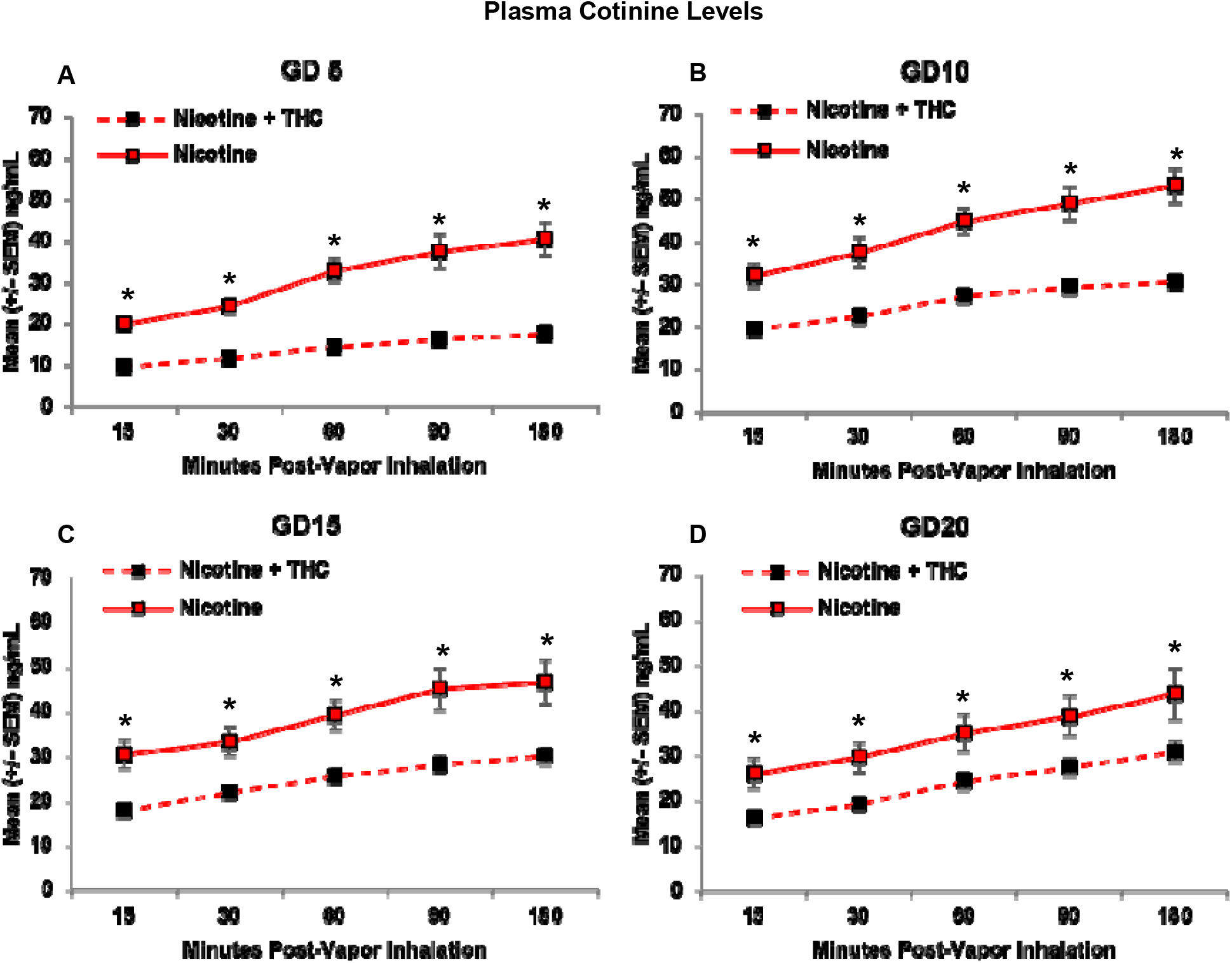
Pregnant dams exposed to the combination of Nicotine+THC had lower plasma cotinine levels than those exposed to Nicotine alone on each day and time throughout the vapor inhalation procedure (A-D). * = Nicotine+THC < Nicotine, *p* < 0.05.

#### 3.1.4.2 THC and Metabolite Levels

Similarly, combined nicotine exposure reduced plasma THC levels in the pregnant dams compared to those exposed to THC alone (F[1,21] = 17.28, *p* < 0.001). Although THC levels increased over days in both groups, consistent with our previous research (Breit et al., 2020), increased plasma levels rose more in the combined group, producing a Day*Nicotine interaction F[3,63] = 3.19, *p* < 0.05, as well as a Time*Nicotine interaction F[4,84] = 4.46, *p* < 0.01; Figure 7). With this gradual increase, although the combination exposure group had lower overall plasma THC levels on GD 5 (F[1,21] = 4.20, *p* = 0.05), 10 (F[1,21] = 8.19, *p* < 0.01), and 15 (F[1,21] = 20.94, *p* < 0.001), levels caught up and did not differ significantly by GD 20.

**Figure 7.**
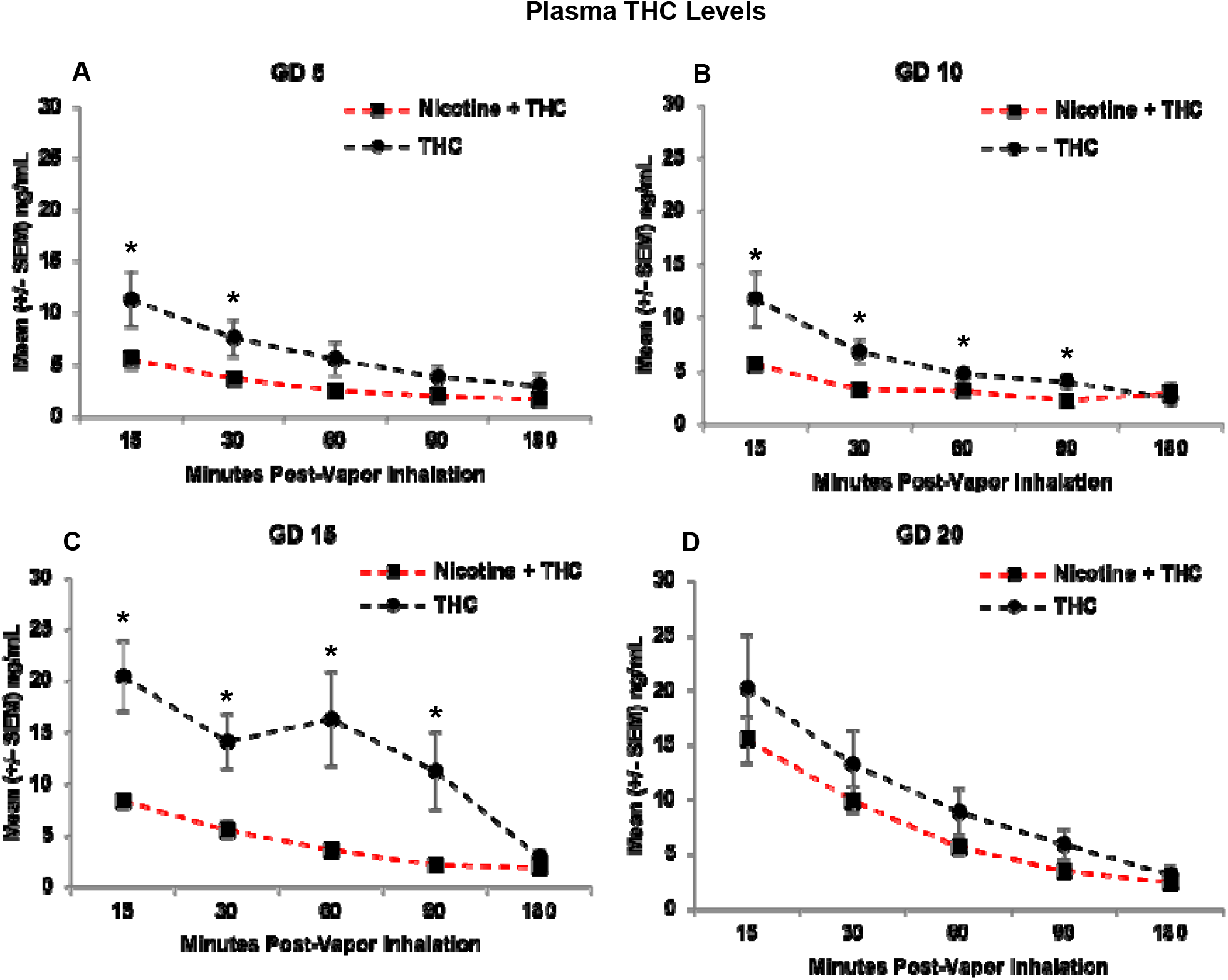
Pregnant dams exposed to the combination of Nicotine+THC had lower plasma THC levels than those exposed to THC alone on gestational days 5 (A), 10 (B), and 15 (C). However, this effect was not seen by the last day of vapor inhalation (D), as plasma THC levels gradually rose across days in both groups. * = Nicotine+THC < Nicotine, *p* < 0.05.

On the first day of vapor inhalation (GD 5), dams exposed to Nicotine + THC had lower plasma THC levels compared to those exposed to THC alone at 15 (F[1,21] = 4.61, *p* < 0.05) and 30 minutes post-vapor inhalation (F[1,21] = 4.73, *p* < 0.05), but not past 60 minutes post-inhalation (Figure 7A). On GD 10 (Figure 7B) and 15 (Figure 7C), the combined exposure group had lower THC levels at all Time points (*p*’s < 0.05) except for 180 minutes post-vapor inhalation. However, by GD 20, the exposure groups no longer differed at any Time point (Figure 7D).

Consistent with THC levels, pregnant dams exposed to the combination of Nicotine+THC also had lower THC-OH metabolite levels than those exposed to THC alone (F[1,21] = 7.87, *p* < 0.05; Figure 8), except on GD 20, producing a Day*Nicotine interaction (F[3,252] = 3.19, *p* < 0.05). Over the Days, THC-OH levels gradually increased among both dams exposed to Nicotine+THC (Day: F[3,33] = 24.43, *p* < 0.001) and THC alone (Day: F[3,30] = 5.17, *p* < 0.05). The combination exposure group had lower metabolite levels on GD 5 (F[1,21] = 4.27, *p* = 0.05), 10 (F[1,21] = 9.14, *p* < 0.01), and 15 (F[1,21] = 13.48, *p* < 0.01), but caught up to THC only group levels on GD 20. In addition, a Time*Nicotine interaction (F[4,84] = 4.88, *p* < 0.01) confirmed that the combined exposure group had lower THC-OH levels at earlier time points (15, 30, 60, and 90 minutes post-vapor inhalation; *p*’s < 0.05), but not toward the end of the sessions.

**Figure 8.**
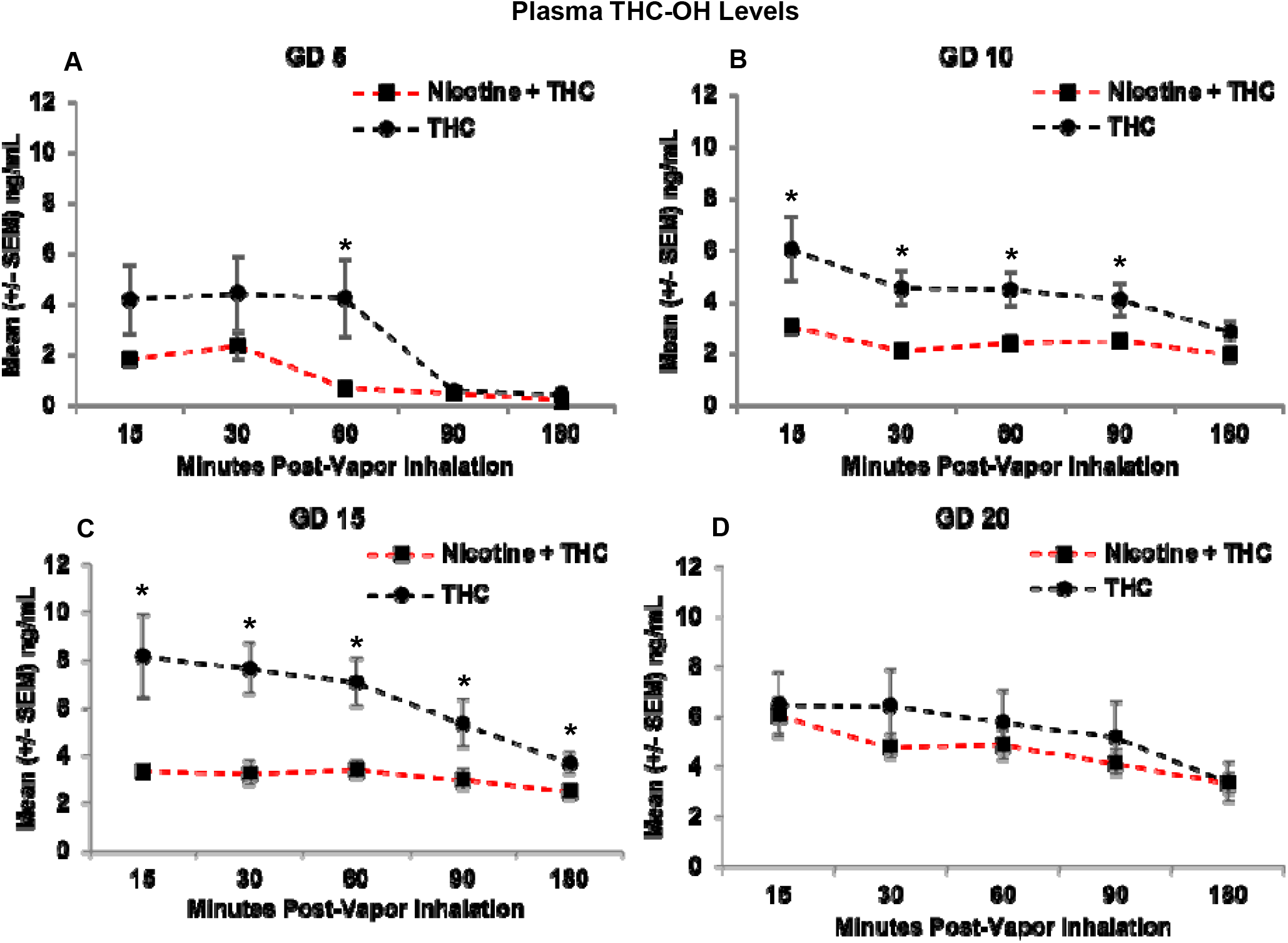
Pregnant dams exposed to the combination of Nicotine+THC had lower THC-OH metabolite levels than those exposed to THC alone on gestational days 5 (A), 10 (B), and 15 (C). However, this effect was not seen by the last day of vapor inhalation (D), as metabolite levels gradually rose in the combination group. * = Nicotine+THC < Nicotine, *p* < 0.05.

Follow-up analyses examined THC metabolite on each individual Day and Time point. On the first day of vapor inhalation (GD 5), dams exposed to Nicotine+THC had lower THC-OH levels compared to those exposed to THC alone only at 60 minutes post-vapor inhalation (F[1,21] = 5.69, *p* < 0.05; Figure 8A). On GD 10, the combined exposure group had lower metabolite levels at all Time points (*p*’s < 0.05) except for 180 minutes post-vapor inhalation (Figure 8B); but by GD 15, this difference was observed at all Time points (*p*’s < 0.05; Figure 8C). Yet, by GD 20, the exposure groups no longer differed at any Time point (Figure 8D).

Similar effects were observed in the THC metabolite levels for THC-COOH; THC-COOH levels were lower in pregnant dams exposed to the combination of Nicotine+THC compared to those exposed to THC alone (F[1,21] = 13.82, *p* < 0.01), but this varied by Day (Day*Nicotine: F[3,63] = 2.69 *p* < 0.05) and Time point (Time*Nicotine: F[4,84] = 2.45, *p* = 0.05; Figure 9). Over the Days, plasma THC-COOH levels gradually increased among dams exposed to Nicotine+THC, but not among those exposed to THC alone. Thus, the combination exposure group had lower overall THC-COOH levels on GD 5 (F[1,21] = 21.52, *p* < 0.001), 10 (F[1,21] = 8.05, *p* < 0.015), and 15 (F[1,21] = 4.86, *p* < 0.05), but not on GD 20.

**Figure 9.**
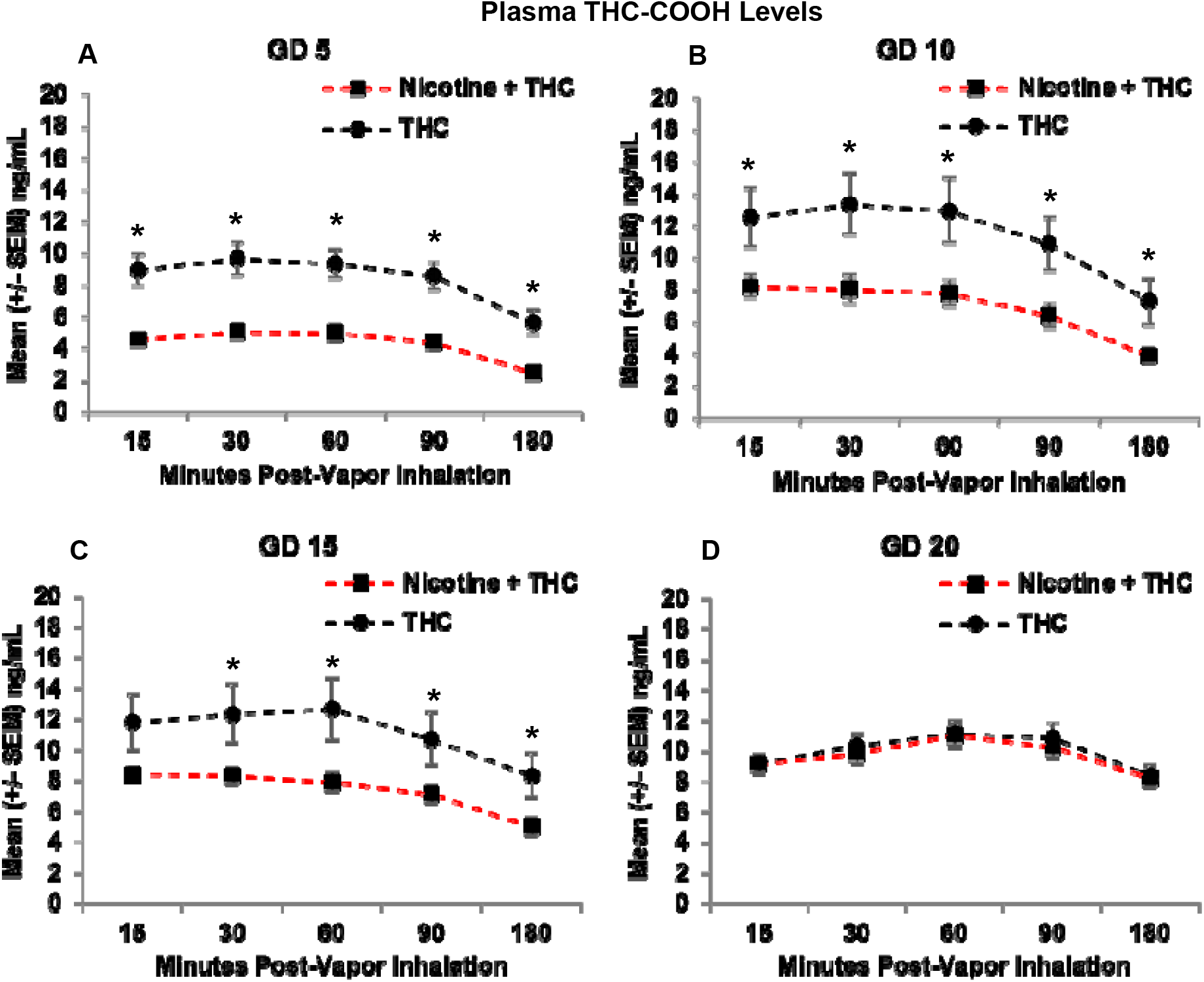
Pregnant dams exposed to the combination of Nicotine+THC had lower THC-COOH metabolite levels than those exposed to THC alone on gestational days 5 (A), 10 (B), and 15 (C). However, this effect was not seen by the last day of vapor inhalation (D), as metabolite levels gradually rose in both exposure groups. * = Nicotine+THC < Nicotine, *p* < 0.05.

Follow-up analyses indicated that dams exposed to Nicotine+THC had lower THC-COOH levels at each Time point on GD 5 (*p*’s < 0.01; Figure 9A) and GD 10 (*p*’s < 0.05; Figure 9B). On GD 15, the combined exposure group had lower THC-COOH levels at all Time points (*p*’s < 0.05) except for the earliest Time (15; *p*’s < 0.05; Figure 9C). By the last day of vapor inhalation (GD 20), the exposure groups no longer differed at any Time point (Figure 9D).

### 3.1.5 Litter Outcomes

Despite the effects on core body temperature and blood levels, prenatal exposure to nicotine, THC, or the combination did not alter basic litter outcomes. No significant differences were observed among gestational lengths, the number of pups born, the sex ratio of litters (Female:Male), or average pup weights at birth (Table 1). Similarly, no differences were observed in the developmental milestone of eye opening (Table 1).

**Table 1.**
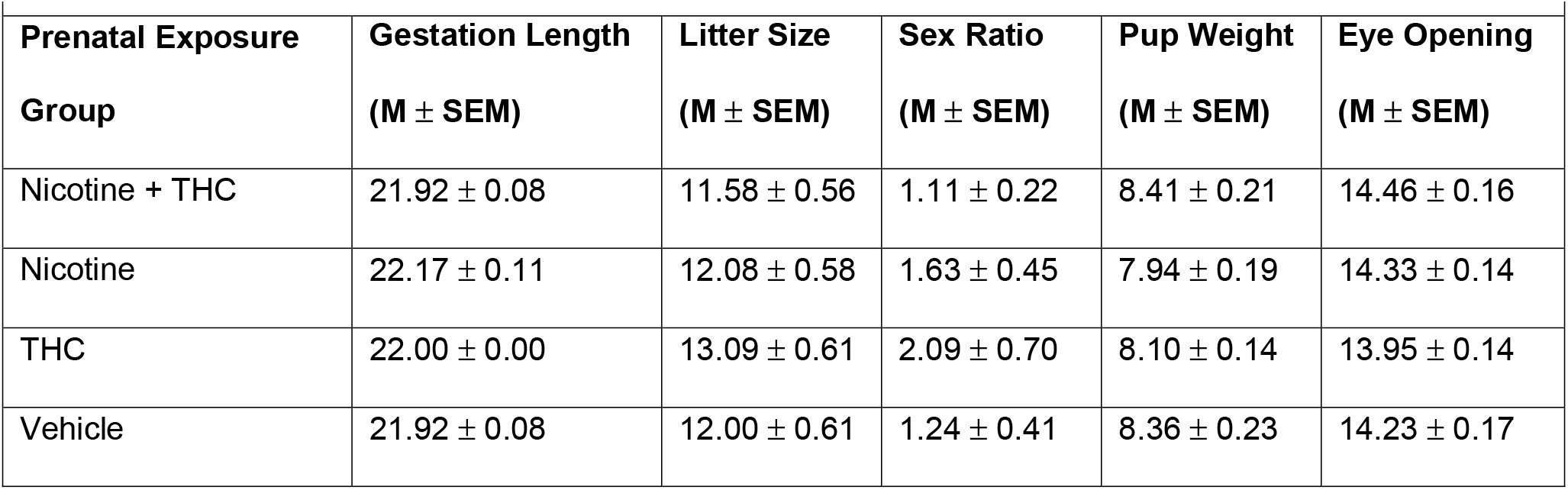
Litter Outcomes Neither prenatal exposure to nicotine nor THC altered basic litter outcomes or the first day of full eye opening.

## 4.1 DISCUSSION

The current study sought to establish a co-exposure model of nicotine and THC via e-cigarette vapor inhalation that can be used safely in pregnant rats. We exposed pregnant subjects to nicotine and/or THC via e-cigarettes daily from GD 5-20, mimicking the human first and second trimester states when drug consumption is most prevalent (Volkow et al., 2019). We successfully demonstrated that this model induces clinically relevant blood drug levels among pregnant rats while avoiding nutritional confounds and poor litter outcomes. To our knowledge, this is the first model to mimic clinically relevant polydrug consumption of nicotine and THC using e-cigarettes among pregnant women in a rodent model, the first to thoroughly monitor multiple maternal and fetal outcomes throughout the exposure period, and the first to demonstrate pharmacokinetic interactions of nicotine and THC in blood levels of pregnant subjects.

In our model, we did not find any changes in maternal body weight growth, daily food intake, or daily water intake resulting from prenatal nicotine or THC exposure throughout the vapor inhalation period. The daily growth and intake of pregnant women consuming either of these drugs is not typically measured, although both prenatal nicotine and THC exposure have been linked to low birth weights (Ernst et al., 2001; Gatzke-Kopp & Beauchaine, 2007; Huizink, 2014). We have previously shown that prenatal THC exposure via e-cigarettes at the same dose used in this study (100 mg/mL) did not alter maternal weight gain, food intake, or water intake (Breit et al., 2020); the other limited studies examining prenatal THC exposure have not reported these measures. Previous research has shown that chronic cigarette smoke in non-pregnant rats reduces body weight, as well as food and water intake (Wager-Srdar et al., 1984), but these outcomes have not yet been measured in pregnant rats, or when e-cigarette exposure is used. These data suggest that at least at the doses used in the present study, that offspring outcome effects (Hussain et al., 2021; Rodriguez et al., 2021), which will be published shortly, are not related to nutritional confounds, nor are pair-fed controls required (Abel & Dintcheff, 1978).

Several studies have previously established that THC exposure via e-cigarettes decreases core body temperatures in non-pregnant (Javadi-Paydar et al., 2018) and pregnant female rats (Breit et al., 2020), similar to the results found in the current study. In contrast, prenatal nicotine exposure reduced baseline core body temperature. It is unknown whether this same phenomenon is observed in pregnant women, although nicotine exposure is known to decrease skin conductance and temperature in humans (Benowitz et al., 1982; Bounameaux et al., 1988; Rowe et al., 1980; Waeber et al., 1984). In non-pregnant rodents, nicotine has been shown to decrease core body temperatures following intoxication via injections (Rezvani & Levin, 2004) and e-cigarette vapor inhalation (Javadi-Paydar et al., 2019). In contrast, we did not find decreased temperatures following nicotine intoxication in the current study, but rather found decreased basal temperatures only following chronic exposure. Notably, pregnant rodents typically show a natural decrease in body temperature as pregnancy progresses (Melanie et al., 1988), although we did not see that among our control subjects.

In addition to the physiological effects on temperature, the blood levels of nicotine, THC, and their metabolites suggest that the doses of nicotine (36 mg/mL) and THC (100 mg/mL) used in our model are clinically relevant. The peak levels of nicotine in our maternal blood samples at this dose replicate those seen in humans and rats given moderate-high doses via various administrative routes, including e-cigarettes (Farsalinos & Polosa, 2014; Farsalinos et al., 2015; Matta et al., 2007; Montanari et al., 2020). The peak levels of blood THC reached in this study are representative of those from low-moderate THC level products, given that high level THC products lead to blood THC levels around 100-150 ng/mL in humans (Andrenyak et al., 2017) and rats (Nguyen et al., 2016). It is, however, important to note that our peak levels were observed 15 minutes post-vapor inhalation, following a 10-minute clearance time before subjects were removed from the chambers. Thus, blood levels were more representative of 25-30 minutes-post drug exposure, and we may have missed the true peak of blood levels of both drugs. Interestingly, THC and its metabolites accumulated in maternal blood over days, which is clinically relevant (Sharma et al., 2012) and has been shown previously in both non-pregnant (Fleischman et al., 1975) and pregnant rats (Breit et al., 2020).

The most striking finding was the pharmacokinetic interactions of nicotine and THC in the blood drug and metabolite levels. Pregnant dams exposed to the combination of nicotine and THC had lower plasma levels of nicotine, THC, and their metabolites compared to those exposed to only nicotine or only THC via e-cigarettes; this effect mostly subsided by the end of the pregnancy (GD 20), since drug and metabolite levels in the dams exposed to the combination approached the levels in those exposed to only one drug, following chronic exposure. This is the first study to co-expose nicotine and THC via e-cigarettes to pregnant rats, and preclinical studies examining prenatal drug exposure rarely report maternal drug blood levels; thus, there is no previous research to compare these findings too.

However, there is limited clinical research on co-consumption in non-pregnant individuals that may be applicable. For example, one study illustrated that users who smoked “blunts” (cannabis rolled in tobacco leaves) had significantly lower plasma THC levels compared to those who smoked “joints” (cannabis wrapped in cigarette paper) at equivalent concentrations, particularly among female subjects (Cooper & Haney, 2009). In this study, both “blunt” and “joint” use yielded similar increases in heart rate and carbon monoxide levels. It was hypothesized that this difference was due to the construction of the “blunts” versus the “joints” since subjects may not be able to reach desired intoxication as quickly with blunts (Cooper & Haney, 2009); however, in the current study, both drugs were administered in the same e-cigarette tank simultaneously, eliminating this possibility. In contrast, other clinical research has shown that transdermal nicotine patch use while consuming cannabis cigarettes enhanced heart rate and reported levels of intoxication more than cannabis alone (Penetar et al., 2005). Similarly, nicotine has been shown to exacerbate several behavioral effects of THC in rodents (Balerio et al., 2006; Valjent et al., 2002). Co-administration of nicotine and THC has been shown to enhance c-Fos expression in various brain regions of rodents (Valjent et al., 2002), and nicotine also potentiates the discriminative effects of THC in rodents in conditioned place preference paradigms (Solinas et al., 2007).

We do know that both nicotine and THC induce the cytochrome P450 (CYP1A2) enzyme, with an additive induction effect following co-use; drugs metabolized by this enzyme may have faster systemic clearance following such enzyme induction (Anderson & Chan, 2016). Separately, both nicotine and THC use in combination with other substances have been shown to decrease plasma concentrations of these drugs in humans (see review by Anderson & Chan, 2016). However, the exact origin of the interaction observed in the current study is unclear and will require further investigation. Regardless, co-exposure to nicotine and THC via e-cigarettes altered each drug’s metabolism, which could have critical implications on fetal development.

Despite the physiological and pharmacokinetic effects observed, we found no alterations in basic litter outcomes. There were no pregnancy or birth complications following prenatal nicotine or THC exposure. Similarly, we found no alterations in gestational length, litter size, average offspring weight, the sex ratio of the litter, or the early developmental milestone of eye opening. Although this is consistent with our previous work looking at prenatal alcohol and THC exposure via e-cigarettes (Breit et al., 2020), it is inconsistent with some clinical research. Prenatal nicotine exposure has been linked to an increased risk of miscarriage, sudden infant death syndrome (Duncan et al., 2009), low birth weight (Ernst et al., 2001; Gatzke-Kopp & Beauchaine, 2007), and other early-life health complications (Bruin et al., 2010); these deficits have also been found following prenatal e-cigarette exposure (Cardenas et al., 2019). Although much less understood, prenatal cannabis exposure has been associated with low birth weight (Huizink, 2014). However, there are many methodological differences between these studies, including (but not limited to) dose levels, routes of administration, and timing of exposure. Nonetheless, these results suggest that this co-exposure model of nicotine and THC exposure via e-cigarettes in pregnant rats can be used to induce physiological changes associated with nicotine and THC consumption while avoiding critical nutritional confounds and alterations in basic litter outcomes.

There are several limitations of this study to address. First, these results only apply to the single doses of nicotine and THC used, which represent moderate-high levels of nicotine consumption and low-moderate levels of THC consumption. Similarly, these results may not generalize to other cannabis products. We chose to initially examine effects of the primary psychoactive constituent, THC. However, cannabis contains more than 500 chemical compounds, including over 100 naturally-occurring cannabinoids, such as cannabidiol (Radwan et al., 2017). Lastly, we acknowledge that the control group in this study was exposed to propylene glycol via e-cigarettes. We did compare data from the vehicle-exposed dams to a small pilot of non-handled, non-exposed dams and found no differences in body weight, food or water intake, gestation length, or any other litter outcome variables. Thus, we used the more appropriate controls for this study, but acknowledge that future research should consider that exposure to e-cigarette vehicles could affect fetal development (Strongin, 2019).

In sum, we have established a novel co-exposure model of nicotine and THC via e-cigarette vapor inhalation for use in pregnant rats. This model induced physiologically relevant effects of nicotine and THC exposure and produced clinically relevant pharmacokinetic interactions in maternal blood drug and metabolite levels. Importantly, these effects were achieved while avoiding effects on nutritional intake, maternal growth, gestational length, and litter outcomes related to maternal and fetal health. With this model, we are currently examining how combined exposure to nicotine and THC via e-cigarettes during gestation alters brain and behavioral development among offspring. These data will provide desperately needed information on the risks associated with e-cigarette use among pregnant women.

## ACKNOWLEDGMENTS

This work was supported by Tobacco-Related Disease Research Program grant (28IP-0026) to Dr. Thomas. It was also supported by a NIAAA training grant (T32AA007456-38) and a NIH Loan Repayment Program award to Dr. Breit. This work was also supported by a Creative Activity and Research Experience Stipend from West Chester University to Mikayla Zeigler. THC was obtained from the NIDA Drug Supply Program. Special thanks to Maury Cole at La Jolla Alcohol Research, Inc. (San Diego, CA) for building the vapor inhalation equipment and for co-exposure advisement. We also want to recognize Dr. Michael Taffe for his aid and advisement in the implementation of this vapor inhalation paradigm, as well as to Dr. Jacques Nguyen, Sophia Vandewater, and Kevin Creehan for training Dr. Breit on intravenous catheter implantation and blood collection procedures. Lastly, we want to recognize the member of the Center for Behavioral Teratology at San Diego State University for assisting in data collection and interpretation, particularly the instrumental efforts of Annie Lei, Brandonn Zamudio, Jaclyn Hanson, and Antonio Ruiz.

